# Simple high-throughput encoding of deep mutational scanning libraries by oligo-based Golden Gate assembly

**DOI:** 10.1101/2025.07.16.665225

**Authors:** Kyrin R. Hanning, Emma J. Walker, Kevin Beijerling, Edward B. Irvine, J. Jordan Steel, William Kelton

## Abstract

Control over mutational library diversity is an essential consideration when engineering proteins, but is often fraught with trade-offs between diversity, specificity, and affordability. Contemporary library assembly approaches often incorporate oligonucleotide pool synthesis to achieve affordable, precise mutagenesis; however, these oligos are often reliant on complex designs to facilitate downstream PCR and/or restriction digests. Direct hybridisation of oligo pools is an overlooked strategy to simplify mutagenesis, especially when paired with a type IIS restriction cloning approach. We validate this approach by designing, hybridising, and deep sequencing single and dual substitution CDR region parts derived from nanobody GA10. Assembly of these parts into a full-length nanobody CDS facilitated the phage display of variant libraries for affinity maturation against its cyclic peptide target. Variants identified through enrichment analysis were expressed in isolation and yielded improved affinities by more than 100-fold. Recent advances in machine learning have successfully inferred improved variants outside of screened library space, but require controlled, multi-mutant libraries. The library assembly approach outlined in this research is well-suited for such approaches.

## Introduction

Engineering proteins via directed evolution has been instrumental for developing enzymes and antibody molecules as drugs and diagnostics. In these processes, point mutations are created within protein-coding sequences and linked via a protein display system (Starr et al., 2020) to protein function, or partitioned (Ma et al., 2018), for functional screening of a desired phenotype. Several methods have become popular for diversity generation within protein sequences, such as the use of error-prone PCR to create distributed mutations throughout a sequence (Cadwell & Joyce, 1992). Challenges with mutational bias and the difficulty of guaranteeing full sequence coverage with error-prone approaches have led to newer methods with synthetic oligo assembly increasing in popularity. Typically, oligonucleotide pools are created, and the second strand is synthesised by PCR prior to cloning into a display vector (Kosuri & Church, 2014). Commercial providers will also create targeted libraries from oligonucleotide pools, but the costs increase drastically with the sequence diversity required. In all cases, a major challenge for directed evolution experiments is covering the enormous potential sequence space of mutations available for screening. For example, the potential combinatorial sequence diversity within a 10 amino-acid antibody CDR region alone exceeds 1 × 10^13^ variants, far surpassing the largest protein engineering libraries created (Bradbury et al., 2011; Plückthun, 2012). Given the size of most proteins being engineered vastly exceeds 100 amino acids in size, new approaches are needed to strategically access protein diversity within the limits of experimental library construction.

One method to systematically tackle this diversity issue is to employ deep mutational scanning (DMS), a successor to alanine scanning that involves the systematic creation of diversity at defined positions within a protein framework (Fowler et al., 2014). In this approach, degenerate codons, such as NNK, are used to saturate mutations at desired locations, or even an entire protein sequence, for screening. Screening of DMS libraries has been widely used as an initial screen to eliminate mutations that are deleterious to protein function and constrain the diversity of subsequent protein engineering libraries (Hanning et al., 2022). However, creating libraries beyond first order DMS (targeting only single sites) to include two or even three fully saturated positions (second or third order DMS) becomes technically challenging and expensive. Higher order libraries are highly desirable to facilitate the capture of epistatic interactions between mutation sites distal in the primary sequence but linked in a three-dimensional folded protein. The information contained within these interactions becomes more important for the training of machine learning models that can assist with protein engineering schemes. Even more desirable are methods that can create a variety of high and low edit distances for robust model training (Taft et al., 2022).

Here, we develop a rapid and cost-effective method for building targeted higher order DMS protein engineering libraries. Degenerate oligonucleotides are designed and assembled by a coiled annealing approach to create overhangs compatible with Golden Gate assembly processes. A directional, multi-part assembly enables the creation of large libraries with targeted diversity for screening in protein display systems. As proof of concept, we apply our method to the affinity maturation of a phage displayed nanobody against a peptide mimetic from the obligate human pathogen *Neisseria gonorrhoeae* and demonstrate several orders of magnitude improvement in affinity. We expect our approach could be used to generate libraries with customised diversity for optimal training of machine learning models and will have utility for other protein engineering applications beyond nanobodies.

## Methods

### Oligos and nanobody sequences

The GA10 nanobody sequence was obtained in-house from a prior screen of a synthetic library (Zimmermann et al., 2020) against PEP-1, a peptide mimetic of the 2C7 lipooligosaccharide epitope from the pathogen *N. gonorrhoeae* (Gulati et al., 1996). The crystal structure of the nanobody used in this work has been submitted to the PDB database (9B0A). All single stranded oligonucleotide pools were ordered as oPools from Integrated DNA Technologies (USA) at a scale of 50 pM per oPool. Wild-type and framework regions were ordered via gene synthesis (Twist Bioscience, USA) for cloning into pUC19 or pAK200-derived vectors using restriction enzyme cloning. All full length nanobody sequences for expression were also ordered from Twist Bioscience.

### Library design

Mutational targeting was designed to encompass both the residues originally defined by the established protocol and additional residues identified by crystallographic analysis as potentially interacting with the antigen, collectively targeting the CDR loops and adjacent residues for diversification. oPool synthesis considerations led to the partitioning of the CDS into five parts, three of which (CDRs 1-3) comprised targeted codons - each effectively mutated via the inclusion of a single or dual NNK codon(s) per designed oligo in a pool. To facilitate scarless CDS assembly of a dual DMS library, each variant oligo incorporated the 5’ four nucleotide overhangs required for correct ligation during Golden Gate assembly. For each CDR part, a total of four oPools were designed and synthesised: single NNK, dual NNK, and their reverse complements, wherein the NNM codon is instead used. Theoretical library diversity is calculated using Eq. 1 where the total number of mutations *M* distributed across *R* parts, where each part *i* has *N*_*i*_ mutable sites, each site mutates in *D* possible ways, and no part exceeds *R*_*max*_ mutations.

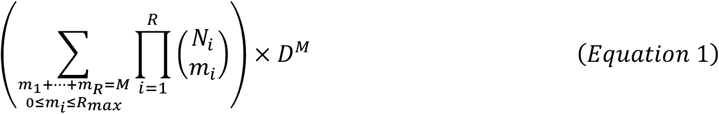

### Oligo pool duplexing and cloning

Lyophilised oPools were resuspended in low TE buffer to 37.5 µM for single NNK pools, or 75 µM for dual NNK pools. Equimolar concentrations of complementary oPools were combined to a total of 300 pmol with T4 Polynucleotide Kinase and supplied buffer (New England Biolabs (NEB), USA) according to manufacturer instructions. The duplexing mixture was then incubated at 37 °C for 40 minutes, and 65 °C for 20 minutes to facilitate 5’ phosphorylation of the ssDNA. Reactions were then immediately heated to 95 °C for five minutes, followed by a cyclic, gradual cooling step. This comprised an initial one-minute incubation at 90 °C, followed by cooling at a constant rate of −0.1 °C·s^-1^ until reaching 65 °C, at which point the reaction would be rapidly heated back to the initial temperature −0.5 °C. This was repeated a total of 50 times, successively reducing the initial temperature to a final of 65 °C before cooling the reaction at −0.1 °C·s^-1^ to 4 °C.

Duplexed oPools were then ligated into pre-prepared plasmid vectors (Supplementary Table 1), generated by BsaI-HFv2 digestion with concurrent quick-CIP (NEB) dephosphorylation, followed by agarose gel excision. Ligations were performed with T4 DNA Ligase (NEB) according to manufacturer instructions and using an estimated 30 fmol insert to 10 fmol plasmid vector. Ligation products were then purified using the QIAquick Nucleotide Removal Kit protocol (Qiagen, Germany) with Monarch® DNA Cleanup Columns (5 µg; NEB). Purified ligation products were transformed into fresh, electrocompetent *E. coli* MC1061 cells via MicroPulser Electroporator (Bio-Rad, USA) according to the manufacturer’s manual, before being entirely plated across several large (140 mm) selective LB agar plates, with ∼100 µL reserved for colony quantification via plated serial dilution. Following overnight incubation at 37 °C, colonies were quantified and resuspended in 1 mL of LB media with 25% glycerol. Purified plasmid was extracted from approximately 2-15 OD600·mL of this culture via QIAprep Spin Miniprep Kit (Qiagen).

### Nanobody CDS assembly

Integration of parts into the full nanobody CDS utilised protocols outlined by New England Biolabs for a Golden Gate assembly strategy utilising BsaI-HFv2 and T4 DNA Ligase (NEB). Briefly, 30 fmol of destination plasmid was combined with 90 fmol of each other plasmid part, BsaI-HFv2, T4 DNA Ligase, and T4 DNA Ligase buffer in a total reaction volume of 50 µL. The reaction was then incubated at 37 °C for five minutes, 16 °C for five minutes for 60 cycles total, terminating with 60 °C for five minutes. Ligation products were then purified using the QIAquick Nucleotide Removal Kit protocol with Monarch® DNA Cleanup Columns (5 µg), and the entire purification was transformed into fresh, electrocompetent NEB 5αF’I^q^ *E. coli* cells. After a brief recovery period (≤one hour; 37 °C; 220 rpm), the entire transformation was plated both undiluted over large (140 mm or 245 mm^2^) selective LB, and as a dilution series on standard plates for quantification. Following overnight incubation at 37 °C, colony numbers were quantified from the diluted plates, and scrape harvested with one mL chilled 2xYT containing glycerol (25% v/v). The OD_600_ of each harvested culture was determined prior to snap freezing for storage at −80 °C.

### Variant library pooling

A total of seven assemblies were performed, each substituting out wild-type CDR part(s) for their equivalent duplexed-oligo-derived NNK part(s). The dual DMS library was generated by proportionally pooling six assemblies; three assemblies simply substituted a CDR for the dual-NNK part to generate the intra-CDR NNK-containing variant sub-libraries; the remaining three assemblies combined the single NNK-containing CDR parts to generate all combinations of the dual inter-CDR variant sub-libraries. In contrast, the high-diversity combinatorial library containing up to six mutations (CMB) was created from only a single assembly of dual-NNK parts for each CDR.

### Phage production and recovery

Variant libraries, thawed from frozen stocks, were inoculated into 50 mL of selective 2xYT medium at an initial OD_600_ of 0.1. The cells were grown at 37 °C to OD_600_ 0.4 - 0.6 before the addition of M13K07 helper phage at an MOI of 20. After resting the cultures for one hour, Kanamycin and IPTG were supplemented (35 µg·mL^-1^, 1 mM, respectively) and growth continued at 30 °C for 8 hours. Phage were harvested by collecting cultures supernatants after centrifugation for 20 minutes at 11,000 xg. Each supernatant was filtered through a 0.22 µm syringe filter and 0.25× the total volume of 20% PEG 8000/2.5 M NaCl was added. The mixture was left on ice for at least 30 minutes and centrifuged at 11,000 xg to pellet the phage before a second round of PEG/NaCl precipitation. All phage were resuspended in 10 mM Tris-EDTA buffer, and phage concentration was determined by plating a serial dilution series of phage incubated with mid-log *E. coli* (OD_600_ ∼0.5) on selective agar plates.

Phage was recovered via infection of 20 mL early-log phase (OD_600_ ∼0.1) NEB 5αF’I^q^ *E. coli* in 2xYT medium supplemented with glucose (2% w/v) incubating at 37 °C. After a one-hour rest period, Chloramphenicol was supplemented (25 µg·mL^-1^) and growth continued at 37 °C until OD_600_ ∼0.3-0.5. Infected cultures were centrifuged at 4,000 xg (4 °C; five minutes) to pellet cells before resuspending them in ∼1 mL of 2xYT containing glycerol (25% v/v) for storage at −80 °C. Aliquots were taken for plasmid extraction and to quantify OD_600_ to facilitate production of the next generation of phage.

### Phage panning

Phage libraries were subjected to a single, non-selective, reinfection round prior to surface panning using a protocol modified from the NEB Ph.D. Phage Display Libraries manual (NEB cat #E8100S) using 96-well ELISA plates. Briefly, coated antigen wells consisted of 50 µL of 10 µg·mL^-1^ neutravidin with 50 nM of the biotinylated 2C7 mimitope (PEP-1) in 50 mM NaHCO_3_, pH 9.5, and were incubated at 4 °C overnight. Subsequent blocking was performed with PBS pH 7.4 w/ 0.05% (v/v) Tween-20 (T), and 2% (w/v) skim milk powder (M) for one hour, shaking at room temperature, followed by six washes with PBST. Phage libraries were diluted to 1 × 10^12^ CFU·mL^-1^ in PBST, with 1 × 10^11^ CFU added per well. Phage were first depleted by incubating for 30 minutes in blocked wells (no coated antigen), before being transferred to coated wells and incubated for one hour (shaking, room temperature). Liquid was then discarded and the plate washed three times with PBST. Bound phage was eluted via incubation with 0.2 M Glycine-HCl (pH 2.2; 1 mg·mL^-1^ BSA) for five minutes before neutralisation with 1M Tris-HCL (pH 9.1) and transfer into a pre-blocked microfuge tube. The entirety of the neutralised phage eluate was subsequently recovered and propagated. Four rounds of selective screening were performed in total. For each round, per replicate, 8 × 10^11^ phage of the dual DMS library, and 1.2 × 10^12^ phage of the combinatorial library were screened.

### Deep sequencing

Purified plasmids of CDR variant parts (cloned, duplexed-oligos), the fully assembled pre-selection libraries (after a non-selective reinfection step to establish growth bias), and the post-selection libraries (after four rounds of screening) were used as templates to prepare amplicons for deep sequencing. Briefly, the variant regions were PCR amplified using custom-designed primers. In a subsequent amplification step, sample-specific barcodes (Illumina Nextera) were introduced to allow pooling of the CDR variant parts and pre-/post-selection libraries into two sequencing runs using (Illumina MiSeq 3; 2×300 bp PE). All sequencing was performed by the Genomics Facility Basel, Switzerland. Quality control and preprocessing of sequencing reads was performed with Fastp (Chen, 2023) and aligned to reference via BWA (Li & Durbin, 2009). Aligned reads underwent a final trimming and polishing step for uniformity with custom Julia scripts (https://github.com/hanningk/MMAtools.jl). Several of these core custom Julia utilities were bundled within a module for downstream analysis.

### Analysis

Aligned, polished reads were classified according to the following criteria: quantity of substitutions within and between the targeted CDR positions, and substitution content. Per position, statistics were determined for substitution frequency and sum unique variants. Enrichment was determined using a simplified Bayesian framework adapted from Bloom (2015). For each variant, the minimum pre-or post-screen count (*x*) determines the sigmoidal pseudo-count prior (Eq. 2) where *a* = 50 is the maximum pseudo-count (applied when *x* ≈ 0), *c* = 1 is the inflection point, and *d* = 1 controls the slope. Given an observed count *n* for the variant in a library size of *N*, its posterior frequency θis drawn (Eq. 3). Fold-change enrichment, *R*, is calculated normalised to the wild-type fold-change (Eq. 4). The reported enrichment score, ⟨log_2_ *R*⟩, is the arithmetic mean of the *M* = 1000 individual log_2_ ratios (Eq. 5).

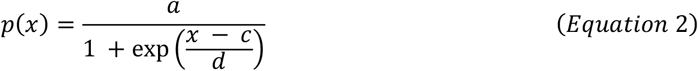

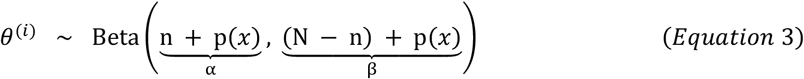

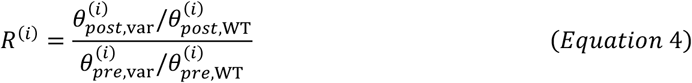

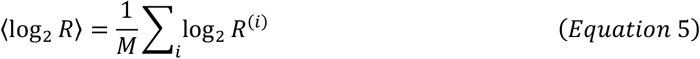

### Nanobody expression and ELISA analysis

Nanobody variants for experimental testing were chosen based on the degree of enrichment and final sequence abundance. Nanobodies were ordered in pET28a vectors for periplasmic expression in BL21(DE3) *E. coli* via the inclusion of a PelB leader sequence. Overnight cultures were used to inoculate 50 mL Terrific Broth media and then grown at 37 °C until an O.D_600_ of 0.6 was reached. Cultures were transferred to 22 °C, induced with 1 mM IPTG, and grown overnight. The next day, the cultures were spun at 4000 xg and the cell pellet resuspended in 5 mL of 20 mM Tris, 150 mM NaCl Tris Buffered Saline (TBS) supplemented with 1 mM MgCl_2_ for periplasmic extraction. The mixture was centrifuged at 11,000 xg for 20 minutes and filtered through 0.22 µm syringe filters. Nanobodies were purified using a 5 mL MabSelect column (Cytiva). Briefly, the column was equilibrated with TBS buffer before loading of the sample. After washing to baseline with TBS, nanobodies were eluted with a gradient from 0-100% over five column volumes (25 mL) of 0.1 M Glycine. The eluted fraction was immediately neutralised with 10% v/v 1 M Tris and buffer exchanged with 10 kDa spin columns back into TBS. Nanobody purity was evaluated by 4-20% Mini Protean TGX Precast gel (Bio-Rad).

To evaluate nanobody binding to the cyclic PEP-1 peptide, ELISA plates were coated with 50 µL volumes of 4 µg·mL^-1^ neutravidin precomplexed with a 1:4 molar ratio of PEP-1 in 50 mM NaHCO_3_. The plates were incubated overnight and blocked with 300 µL of TBS containing 2% skim milk powder w/v for one hour at room temperature. Nanobodies were prepared in TBSM and serially diluted down the plate, left at room temperature to bind for a further hour, and removed by washing three times with TBS containing 0.05% Tween20. Bound nanobodies were detected by incubation for an hour with 50 µL of anti-VHH in TBSM at a 1:10,000 dilution (Clone: 96A3F5, Genscript). The plate was washed 3X in TBST, developed with 50 µL Ultra TMB substrate (ThermoFisher), and quenched with 50 µL of 1M H_2_SO_4._ Absorbance was quantified at 450 nm via a plate reader.

## Results

### Efficient and Controlled Duplex Formation of Mutagenic Oligonucleotides

To create a modular approach for the flexible generation of diversity within targeted regions of protein sequence libraries, we demonstrate a gene assembly method using degenerate oligo hybridisation without PCR amplification. We selected a candidate nanobody with moderate affinity (PDB: 9B0A) raised against a cyclic peptide vaccine candidate (PEP-1) against *Neisseria gonorrhoeae* for affinity maturation via phage display. The nanobody was originally derived from a synthetic nanobody library of complementarity-determining region (CDR) variants (Zimmermann et al., 2020). Using information from the crystal structure, we expanded the set of CDR residues targeted for variation in the original nanobody library to include adjacent sites involved in PEP-1 binding (Fig. 1A). These redefined CDRs represent logical parts for Golden Gate assembly methods, and their lengths are ideal for synthesis as oligonucleotides. The nanobody CDS was split into six contributing parts: three target CDRs and three non-targeted parts, optimising assembly fidelity of the overhangs (Weber et al., 2011). Each CDR part was synthesised as “single” and “dual” substitution oligo pools, within which targeted codons are substituted with single and dual-concurrent NNK degeneracy, respectively. To facilitate PCR-free mutagenesis, the equivalent, reverse complement oligo pool was also synthesised, incorporating NNM degeneracy. Duplexing of the two cognate pools would yield dsDNA with 5’-four nucleotide overhangs – facilitating immediate post-duplexing ligation. To further improve fidelity of this ligation, 5’ phosphorylation was performed on the ssDNA oligos immediately prior to duplex formation (Fig. 1B). All parts were co-assembled with wild-type framework parts to generate full nanobody CDS sequences (Fig. 1C). To establish the flexibility of diversity profiles achieved with the method, we created libraries with tailored high and low diversity profiles of up to 6 (combinatorial) and up to 2 (dual DMS) mutations per variant respectively. All nanobody variants were expressed on M13 bacteriophage and screened against immobilised cyclic peptide antigen for four rounds of selection (Fig. 1 D).

**Figure 1.**
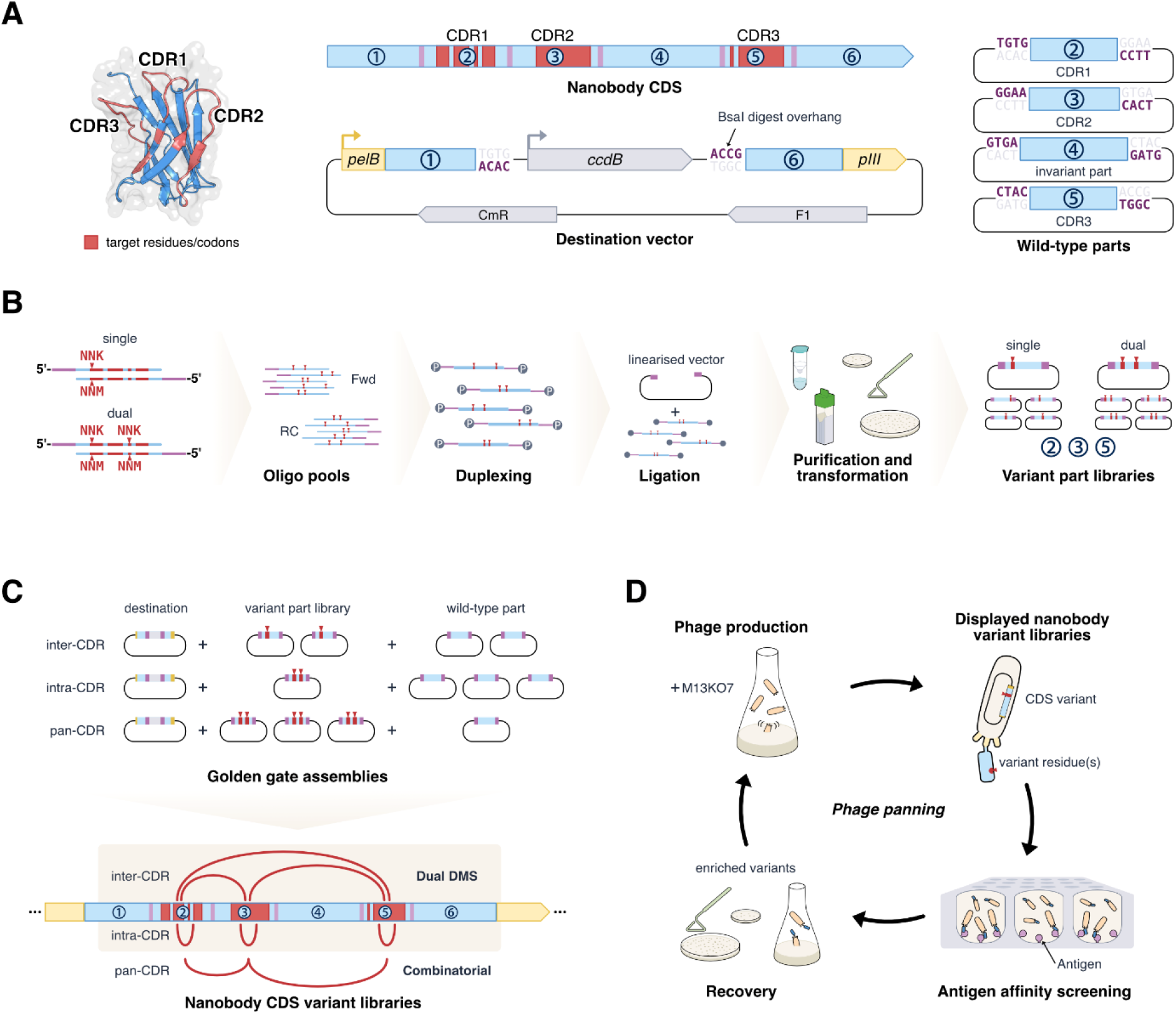
Variant nanobody library design and construction approach. Library design. **(A):** Residues encompassing the CDR regions were targeted for mutation (red). The nanobody CDS was split into six parts to facilitate effective synthesis of the CDRs (parts 2, 3, and 5) as separate oligo pools. Part boundaries were determined by suitability of overhangs (purple/pink) for Golden Gate assembly via BsaI restriction-enzyme digest. The nanobody CDS termini (parts 1 and 6) were synthesised with an internal counter-selection cassette (*ccdB*) flanked by BsaI recognition sites. The fragment was cloned into a phagemid (F1 ori) destination vector, with 5’ *pelB* and 3’ M13 phage *pIII* fusions (yellow) to facilitate phage expression upon assembly with the remaining CDS parts (2-5). These parts were synthesised with flanking BsaI sites and cloned into pUC19-derived vectors prior to Golden Gate assembly. **Variant part generation (B):** Oligos of the three parts representing the CDRs (2, 3, 5) were designed as single or dual NNK-codon containing libraries and synthesised as separate forward (FWD) and reverse complement (RC) oligo pools (oPools). Each oligo contained a unique 5’ Golden Gate overhang, which after 5’ phosphorylation and duplexing via heat-cool cycles, enabled their immediate ligation into overhang-matched pre-digested vectors. The ligation products were immediately purified and electro-transformed to yield separate single and dual variant libraries for each part. **Golden Gate assemblies (C):** By substituting wild-type parts with variant library parts during Golden Gate assembly, seven nanobody CDS variant libraries were generated. Six of these formed the dual DMS library; ≤2 substitutions total, either inter-or intra-CDR. The seventh library was assembled from the dual intra-CDR parts to yield a high diversity combinatorial library; ≤ 6 substitutions total, ≤2 intra-CDR. The assembled libraries were electro-transformed into NEB 5-alpha F’Iq *E. coli*. **Variant screening (D):** All bacterial libraries were cultured and infected with M13KO7 helper phage to produce nanobody-pIII fused phage. This recombinant phage was then screened against an immobilised antigen (PEP-1) before recovery via infection and plating of fresh *E. coli*. This panning process was repeated for multiple rounds.

We first sought to validate the depth of coverage and fidelity of the oligo hybridisation method. Each of the six hybridisations for the single NKK and dual NNK variant pools were independently repeated before cloning into storage plasmids. Colony counts revealed a coverage depth of the theoretical diversity of between 292-and 1,300-fold for the single NNK parts, and between 5.8-and 50-fold for the higher diversity dual NNK parts (Table 1). We observed a reasonable degree of consistency across both replicates. Minimising the time between oligo hybridisation and subsequent ligation was critical to ensuring a high coverage of variant diversity.

**Table 1.** Transformation yields for replicate (#1, #2) oligo pool duplexing. Yield represents the total approximate colonies harvested derived from dilution series, and assumed depth of total, targeted, non-redundant NNK variants.

| Cloned duplex |  | Theoretical Diversity | Yield (depth) #1 | Yield (depth) #2 |
| --- | --- | --- | --- | --- |
| Single NNK | CDR1 | 346 | $3.0 \times 10^5$ (867×) | $1.1 \times 10^5$ (318×) |
| | CDR2 | 408 | $1.2 \times 10^5$ (294×) | $2.7 \times 10^5$ (662×) |
| | CDR3 | 377 | $4.9 \times 10^5$ (1300×) | $1.1 \times 10^5$ (292×) |
| Dual NNK | CDR1 | 54,447 | $2.7 \times 10^6$ (50×) | $9.0 \times 10^5$ (17×) |
| | CDR2 | 76,860 | $4.5 \times 10^5$ (5.8×) | $9.5 \times 10^5$ (12×) |
| | CDR3 | 65,173 | $8.3 \times 10^5$ (13×) | $1.8 \times 10^6$ (28×) |

### Characterisation of Initial Variant Space by Deep Sequencing

To probe coverage of the theoretical diversity at single nucleotide resolution and evaluate positional bias due to hybridisation, we next deep sequenced the individual part libraries. From the resulting paired end reads, the highest quality sequences were processed, aligned, and polished as single end reads (Supplementary Figure S1, S2). For the single site libraries, we observed 100% coverage of the theoretical diversity across all parts sequenced in both replicates. In the dual NNK libraries, we observed >97% coverage of the expected variant codons (>99% when translated). We next interrogated the mutational frequencies at each position within the CDRs (Fig. 2A). As expected, the mean mutational frequency within the dual site assemblies is much lower than in the single site assemblies. Likewise, single site assemblies have fewer unique reads than their dual site counterparts. One notable exception was the CDR2 assembly, which showed a reduced mutational frequency towards the 3’ end of the assembly, indicative of less diversity in this region. The use of deep sequencing also enabled evaluation of background or non-expected sequences that fell outside the design constraints of NKK codons. While most background sequences in non-mutated positions were consistently low, higher frequencies were observed in positions where NNK codons were used. Additionally, no obvious increase in mutational frequency is observed in codons that overlapped the Golden Gate overhang.

**Figure 2.**
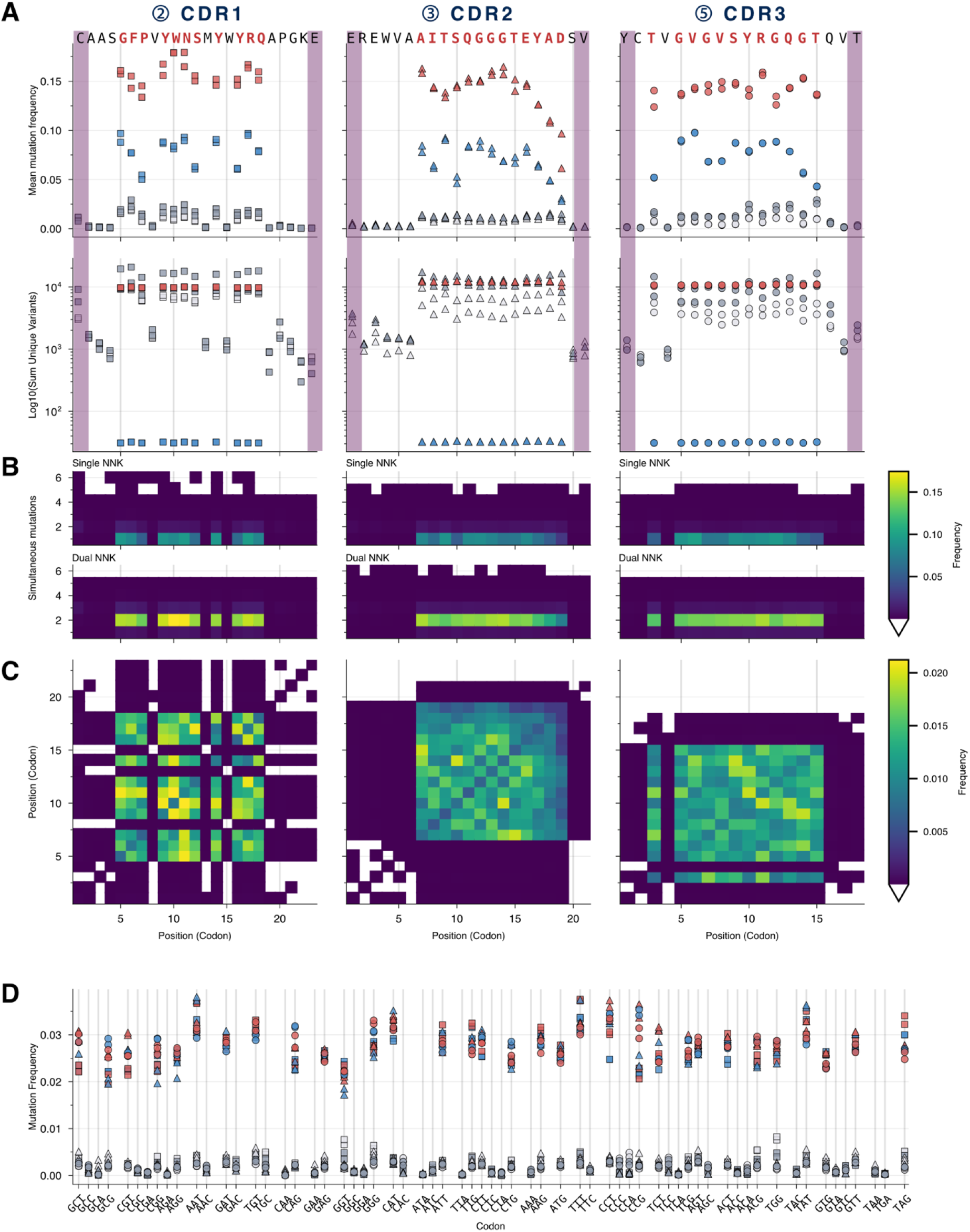
Codon variance on a per position basis within duplexed NNK oligo pools. Parts 2, 3, and 5, corresponding to CDRs 1, 2, and 3, respectively were cloned into storage plasmids and deep sequenced. (**A**) Plot of variant diversity within each part showing codon mutational frequencies and the number of contributing unique variants. Shown in duplicate are variants that met the intended design constraints (including permitted NNK codons, number of mutant codons) for the single (blue) and dual (red) NNK libraries. Also included are variants that fell outside the design criteria for the same single (light grey) and dual (dark grey) NNK libraries. The wild-type amino acid sequence is shown, with targeted residues highlighted in red, and the type IIs restriction site overhang positions shown in pink. (**B**) Heatmap of mutational biases in the single and dual NNK libraries. Sequences were stratified by mutation count, and the frequency at which mutations are observed was calculated at each position within the CDRs. Individual squares represent the mean of two replicates. (**C**) Two-dimensional heatmap of mutational bias within the dual NNK library parts. For variants that passed the design criteria (≤2 codon mutants), bias was evaluated by correlating observed mutation frequency at each position. Each square represents the mean of both replicates. (**D**) Calculated codon preferences of variants meeting/failing the design constraints within the single (blue/light grey) and dual (red/dark grey) libraries across each of the three CDR parts (1: square; 2: triangle; 3: circle).

A global evaluation of mutations within each hybridised part also confirms a strong bias towards positions where diversity is desired (Fig. 2B). Variants falling outside the expected mutational range for each part typically contained more mutations than expected, up to a maximum of six. When the dual site libraries were investigated in greater detail by mapping linkage between mutational positions (Fig. 2C), the CDR2 assembly again presented with a strong bias of low mutational frequencies towards the 3’ end. The CDR1 assembly also displayed a stronger tendency for linked mutations than the other two CDR assemblies. We lastly looked at individual codon frequencies, which were remarkably consistent for single and dual site assemblies suggesting good representation of diversity is maintained at a codon level noting that translation of the sequences to protein will eliminate some redundancy (Fig. 2D).

Golden Gate assembly was used to assemble full-length constructs. Post-assembly transformation yielded a range of library coverage efficiencies (Table 2), while still enabling construction of a large-scale dual DMS library with minimal bottlenecking. Based on colony count, the diversity of the assembled, dual DMS sub-libraries exceeded the theoretical diversity (≥4.7×), whereas the combinatorial library reached ≤1.8 × 10^−8^ % of theoretical maximum.

**Table 2.**
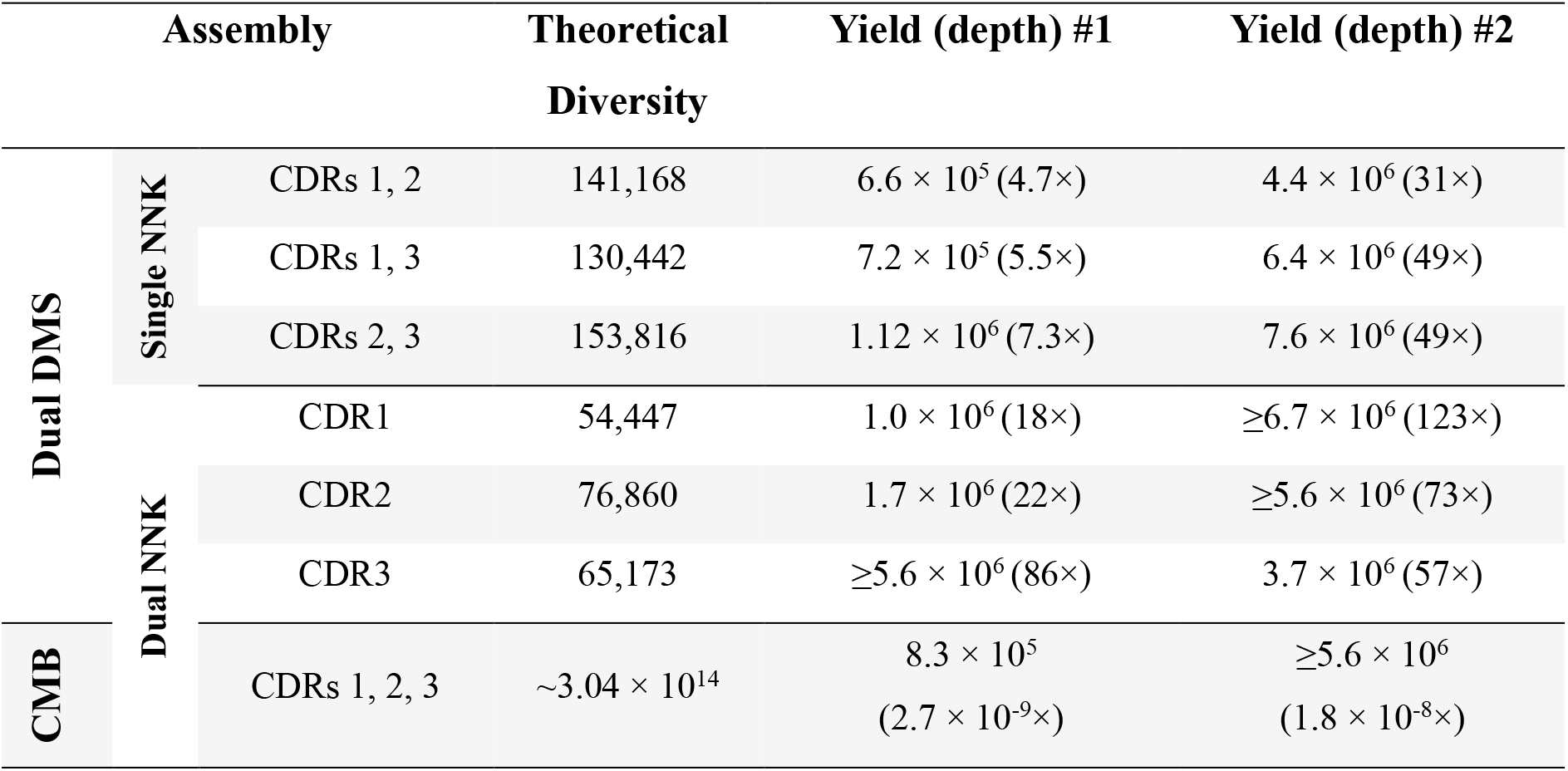
Golden Gate assembly yields for replicate (#1, #2) assemblies contributing to the dual DMS libraries and combinatorial (CMB) libraries. Yield represents the total approximate colonies harvested, derived from dilution series, and assumed depth of total, targeted, and non-redundant NNK variants. Theoretical diversity was calculated for each library using Equation 1.

Deep sequencing of the pre-and post-selection libraries facilitated a discrete analysis of the assembly products, in addition to the intended enrichment analysis. These libraries underwent identical processing, alignment, and polishing as overlapping paired end reads (Suppl. 1, 2).

Dual DMS and combinatorial assemblies largely reflect the targeted per position bias (Fig. 3A, 3B) seen with the CDR parts (Fig. 2A). For both dual DMS and combinatorial libraries, most reads represented variants that conformed to the intended design criteria of the respective libraries (68% of dual DMS and 93% of combinatorial library reads). The larger population of non-conformant variants in the dual DMS libraries was due to a number of variants containing

**Figure 3.**
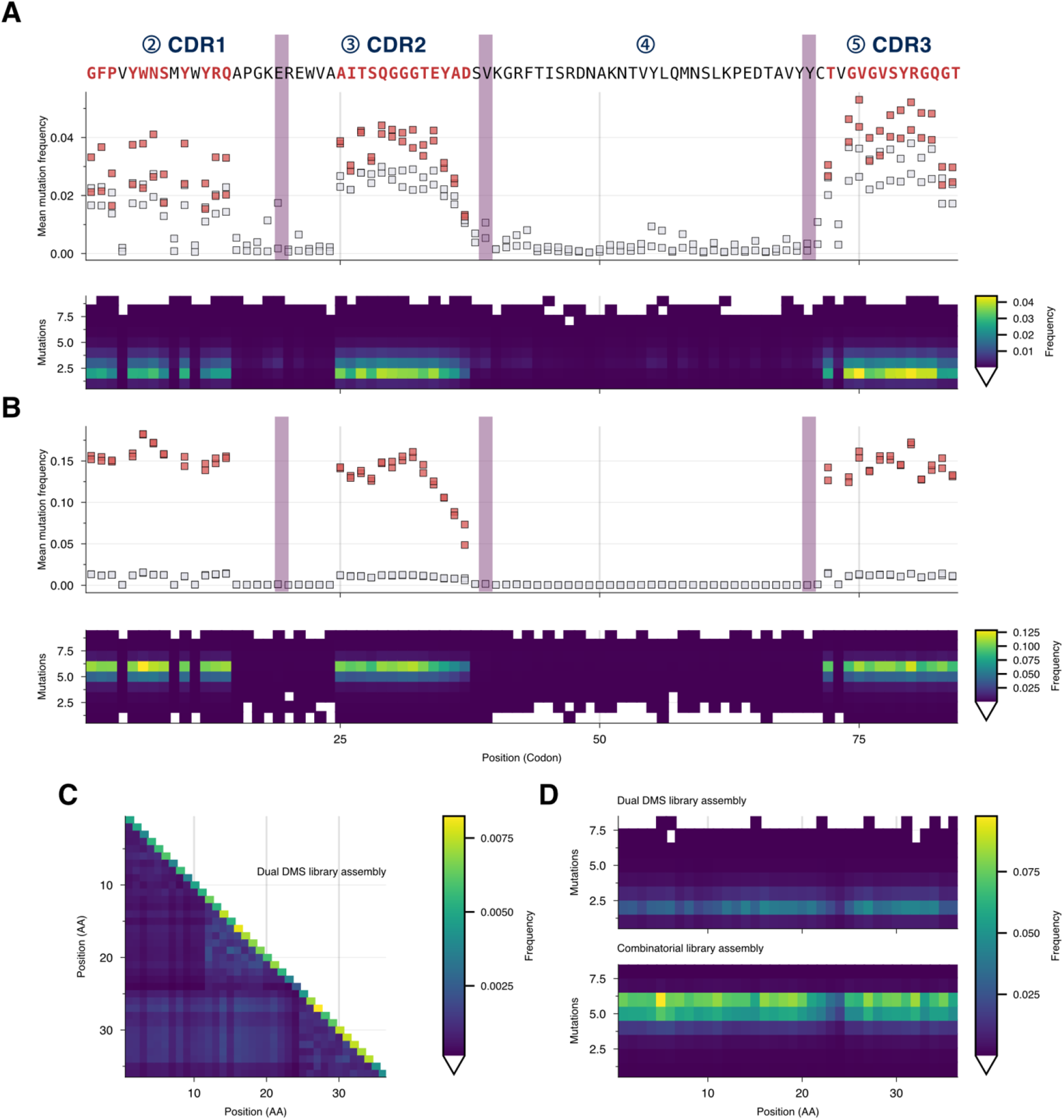
Amino acid variance derived from the sequencing of fully assembled variants from the pre-selection library. **(A)** Plot of variants from the dual DMS library displaying the frequencies at which the observed codons deviate from wild-type (average of two replicates). Shown are variants that fall within the intended design constraints (Red; ≤2 intra-CDR variants; ≤2 inter-CDR variants; no TGA or TAA stop codons) and variants that do not (Grey). The heatmap shows the frequency of observed mutations at each position within dual DMS library variants, stratified by total mutation count. **(B)** Plot of variants from the combinatorial library displaying the frequencies at which the observed codons deviate from wild-type (mean of two replicates). Displayed are variants that fall within the intended design constraints (Red; ≤2 intra-CDR variants; ≤6 inter-CDR variants; no TGA or TAA stop codons) and variants that do not (Grey). The heatmap shows the frequency of observed mutations at each position within combinatorial library variants, stratified by total mutation count. **(C, D)** Variants were translated, and residues targeted for mutation within each of the CDRs were concatenated. **(C)** Two-dimensional heatmap of mutational bias within the dual DMS library assemblies (mean of two replicates). For variants that passed the design criteria (≤2 amino acid changes), bias was evaluated by correlating observed mutation frequencies at each position. **(D)** Heatmap of mutational biases in the dual DMS (top) and combinatorial (bottom) libraries. Sequences were stratified by mutation count, and the frequency at which mutations are observed was calculated at each position within the concatenated CDRs.

> 2 substitutions. Regarding coverage of the theoretical library diversity, the dual DMS libraries achieved 36% (225,170 variants) and 62.5% (387,429 variants) for the two replicates. In contrast, the combinatorial library replicates showed much lower coverage, with 178,909 variants (5.9×10^−8^%) and 952,365 variants (3.13×10^−7^%), respectively.

Due to comparatively low off-target substitution, and to aid with enrichment analysis, reads were condensed into spans of the 36 target residues, comprising CDR1 (1-11), CDR2 (12-24), and CDR3 (25-36). Amongst the single and dual-substitution variants within the dual DMS library, there is a bias towards dual-substitutions with CDR3 residues, and internally within CDR2 (Fig. 3C). Multi-substitution trends within the condensed variants remained comparable to the full-length assembled sequence (Fig. 3D).

The enrichment of each variant was determined as the fold change from wild-type abundance. Typically, DMS analysis is restricted to those variants observed in the pre-screening library, this was enforced for analysis of the dual DMS and combinatorial libraries (Fig. 4A). Here, due to potential under sampling in the combinatorial library, we also considered the situation where variants were observed in the post-screened library but not in the pre-screened library. This approach captures a wider sequence space at the risk of reducing accuracy of enrichment prediction. For the dual DMS library replicates, both approaches showed a high degree of enrichment convergence, with Pearson correlation values of 0.84 and 0.83 for the restrictive and relaxed approaches. Deeper analysis of replicate sequence similarity in pre-screened libraries revealed a more modest Jaccard index (fraction of shared sequences) of 0.18 and 0.24 for the restrictive and relaxed approaches across the two replicates. For the combinatorial library replicates, under-sampling of the potential library diversity meant a narrow pool of variants were observed in each of the replicates. The restrictive analysis approach only analysed a pool of 999 unique variants, which was expanded to 44,173 when using the relaxed approach. Overall, the Jaccard indices were < 0.01 irrespective of the analysis method between these replicates.

**Figure 4.**
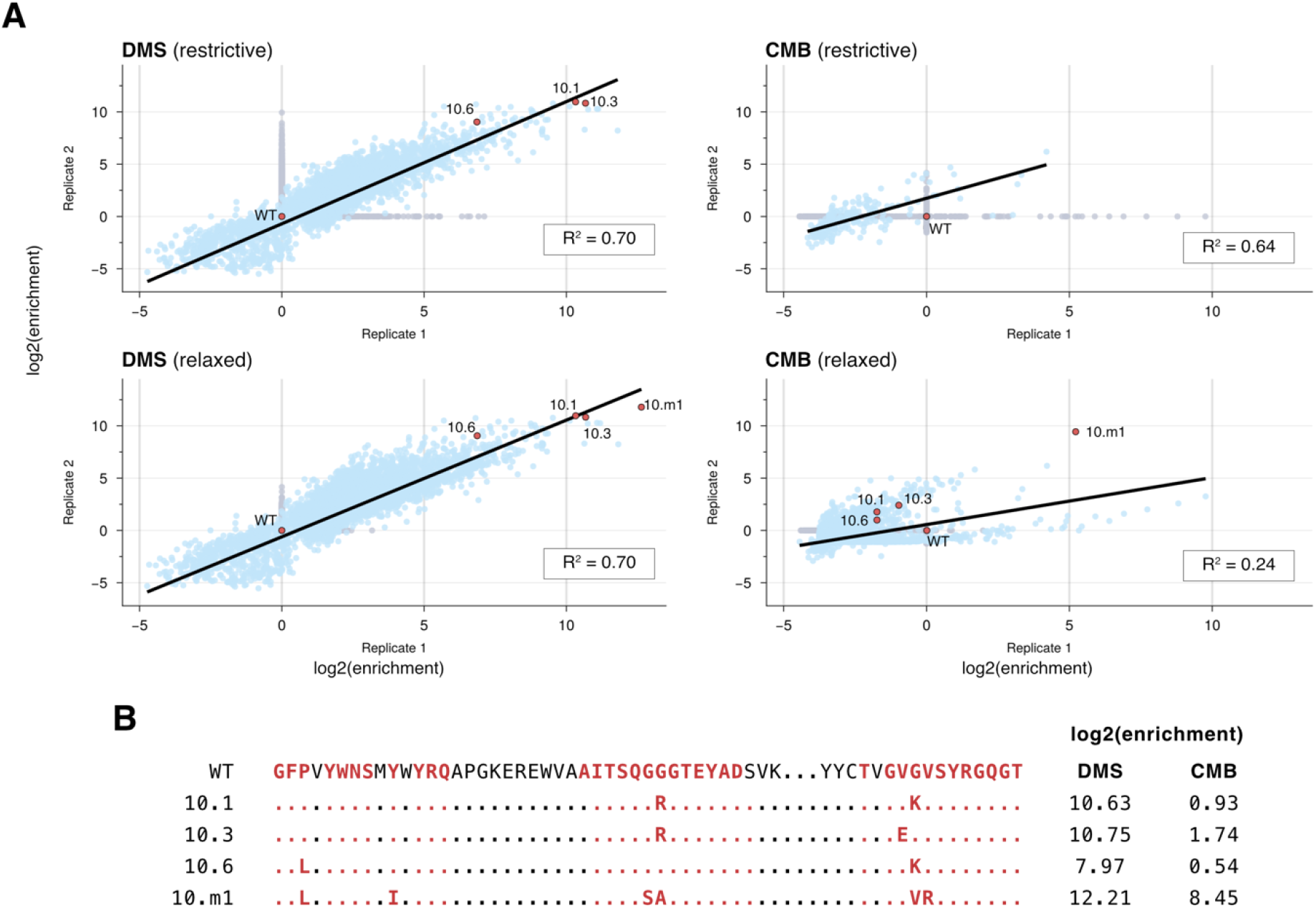
Summary of dual DMS and combinatorial enrichment. **(A)** Distribution of replicate enrichment ratios for the DMS (left) and combinatorial (right) libraries. Enrichment ratios were calculated by taking the mean posterior log_2_ enrichment relative to wild-type (WT). Unique sequences observed in both replicates are coloured blue, whereas variants only observed in a single replicate are in grey. A linear regression was also fitted to shared replicate data using ordinary least squares (solid line) and the associated R^2^ values reported. **(B)** Sequences of nanobody variants selected for expression. Each variant is represented as a red point on the enrichment plots.

To determine whether nanobody variants represented with high enrichment scores possess higher affinity, we selected a small panel of variants for experimental affinity evaluation (Fig. 4B). These variants were chosen based on high abundance in the raw read counts and high ranking in enrichment scores across the two dual DMS replicates. Despite under sampling, we also selected one variant with six unique mutations (10.m1) that appeared in all libraries that had significantly higher raw read counts and enrichment than any other variant.

Nanobody variants were expressed in the periplasm of *E. coli* and purified using a conformationally selective protein A-based resin. SDS-PAGE analysis showed nanobodies at the expected sizes, noting that the WT variant retains additional C-terminal sequences as a result of cloning from the original synthetic library (Fig. 5A). Variants were evaluated for binding to the cyclic PEP-1 peptide relative to the wild-type nanobody. From the dual DMS library, the 10.1 and 10.3 nanobody clones displayed similar binding and showed improvements over the wild-type variant of more than 100-fold (Fig. 5B). The third ranked clone, 10.6, also displayed improved affinity with a more than 50-fold increase relative to wild-type. A single variant with six mutations was chosen from the combinatorial library for analysis. Surprisingly, and despite a high enrichment score, the 10.m1 variant had a more modest but still significant affinity increase over WT of more than 20-fold. Nonetheless, all selected variants displayed large improvements over the base nanobody sequence, highlighting the utility of this approach for rapid nanobody affinity maturation.

**Figure 5.**
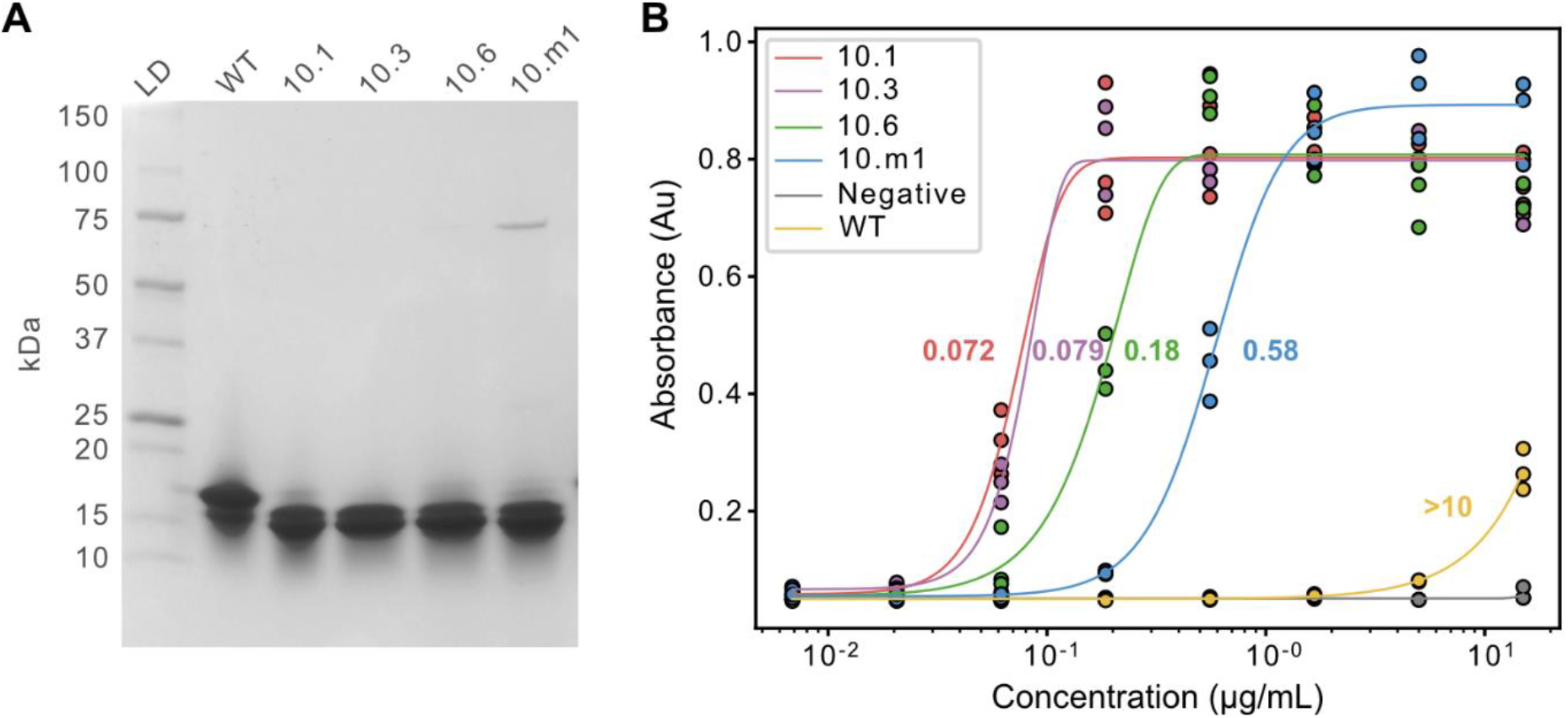
ELISA analysis of engineered nanobody variant affinity. (**A**) Nanobody variants were selected with the highest-ranking enrichment scores from both the dual site (3 variants) and combinatorial libraries (1 variant). Nanobodies were expressed in *E. coli*, purified with MabSelect™ PrismA resin, and run by SDS-PAGE using a 4-20% gradient gel. The wild-type variant was expressed in parallel. (**B**) Nanobody affinity was measured by ELISA against PEP-1 peptide coated plates. Nanobodies were diluted from a starting concentration of 15 μg/mL, with the anti-EGFR nanobody 7D12 used as a negative control. EC50 values were obtained by fitting of 5-parameter logistic (10.1, 10.6, 10.m1) or 4-parameter logistic models (WT).

## Discussion

Synthetic biology approaches such as Golden Gate cloning are providing increased flexibility for protein engineers when designing libraries. In this study, we developed a method to create protein libraries with targeted and precisely controllable diversity profiles within larger protein sequences. We applied this method to create high and low edit distance libraries for the affinity maturation of a candidate nanobody with moderate affinity against a peptide vaccine candidate. Deep sequencing was used to characterise each step of the library construction process, revealing a minimal degree of codon bias and high coverage of the theoretical diversity at the individual part level. The final assembled nanobody libraries were panned against the cognate peptide, resulting in binders with greater than 100-fold improvements in affinity.

Unlike traditional library construction approaches, in which error prone methods are employed (Lee et al., 2018), targeted diversity allows a more constrained library focused on regions most likely to deliver a desired phenotype. This focusing becomes increasingly important as the size of the protein being engineered increases. More recent approaches using gene synthesis in larger oligonucleotide pools have greatly improved the ease of targeted library construction containing saturation mutagenesis, but lack flexibility in the control of edit distances (Fowler et al., 2014; Melnikov et al., 2014). However, obtaining large library sizes remains challenging and many of these methods use PCR to amplify regions of diversity, which increases the risk of variant bias/drop-out before screening. A key difference in our approach has been to use complementary oligonucleotide pools for direct assembly with overhangs ready for Golden Gate Cloning. While we first cloned these pools into intermediate vectors to evaluate hybridisation fidelity by deep sequencing, direct assembly without prior cloning may further improve the speed of library generation. Access to alternative deep sequencing techniques, such as Oxford Nanopore, may reduce screening timelines while enabling an entirely PCR-free solution to data collection.

During oligonucleotide assembly, deep sequencing led us to notice a drop in mutation frequency towards the 3’ end of the CDR2 parts in both the single and dual site assemblies. The CDR2 part possesses Golden Gate overhang sequences closest to the positions of introduced NNK diversity (3 bases). Depending on the GC content of the sequence, these results suggest increasing the distance between the proximal NNK codon and the Golden Gate overhang to at least five bases to improve oligo pool annealing. The use of deep sequencing further allowed us to evaluate the appearance of codons falling outside NNK degeneracy design profiles. It was surprising to note that background mutations were low in positions not targeted for variation but more concentrated where NNK codons were present. We speculate these background rates could be derived from mismatch repair processes occurring when annealed oligos do not find perfectly complementary sequences (Khilko et al., 2018; Kosuri & Church, 2014), rather than stemming from sequencing or oligo synthesis errors, which would distribute errors more evenly across the sequence. However, given codon degeneracy during translation, and the ability to bioinformatically filter out these variants, they pose little issue when selecting for enriched variants.

In this work, the highest affinity nanobody variants were retrieved from the dual DMS libraries with up to two mutations. Under-sampling in the combinatorial library replicates resulted in comparison of enrichment scores across a very limited pool of shared variants. For high diversity libraries, this means replicate analysis is unlikely to add much additional accuracy over the screening of a single library. Further, our finding that the 10.m1 variant with six mutations from wild-type did not have improved affinity over the variants selected from the dual DMS libraries also highlights another potential challenge with phage display methods, whereby affinity and expression improvements are challenging to decouple.

While single site DMS is a relatively accessible method, there are few examples of higher order DMS owing to the challenge of library construction. Accessing this diversity will enable easier systematic profiling of epistatic interactions within proteins. Alongside epistasis, the ability to control the depth of diversity within select regions of proteins has particular promise in machine learning guided protein engineering. These methods allow extrapolation of the binding landscape well beyond the experimental sequence space screened. Evidence is emerging that the edit distance distribution of libraries is a significant contributor to the accuracy of deep learning models predicting protein-protein binding interactions (Erlach et al., 2025; Frei et al., 2025; Taft et al., 2022). The modular system developed here allows diversity to be controlled by the mix of individual parts combined to form final library assemblies. Further strategies, such as the incorporation of triple site NNK parts and the intentional doping of wild-type parts, could provide further control when creating libraries. Further, noise in the form of false positive and false negative binders is a particular challenge during model training. Although approaches to mitigate noise with small subsets of high-quality data have emerged (Minot & Reddy, 2024), the ability to eliminate sources of noise using deep sequenced replicates may assist with model accuracy.

Control over protein library diversity has historically been challenging. Here, we present a PCR-free library building approach using short oligonucleotides that is both affordable and accessible to laboratories with molecular biology skills. Furthermore, we expect the modularity and scalability of the method to be broadly applicable beyond protein display and engineering applications.

## Acknowledgements

Consumables for this work were supported by the Waikato Medical Research Foundation (Grant #320). K.R.H, E.J.W, and K.B were supported by Waikato doctoral scholarships. E.B.I was supported by an ETH postdoctoral scholarship. W.K was supported by MBIE Smart Ideas (Grant: UOWX2307) and Marsden Fund grants (Grant: MFP-UOW2301).

## Author contributions

K.R.H., and W.K. conceived of and designed the study. K.R.H., E.J.W., K.B, J.J.S, and W.K. performed laboratory experiments and created reagents used in the study. K.R.H., W.K. and

E.B.I. performed the Illumina sequencing. K.R.H undertook the bioinformatic analysis. K.R.H., and W.K. wrote the manuscript.

## Notes

### Competing Interest Statement

The authors have declared no competing interest.

