## Supplemental material for "Simple high-throughput encoding of deep mutational scanning libraries by oligo-based Golden Gate assembly"

**Supplementary Material**

### Supplementary Tables

**Table S1. Oligo/plasmid sequences used in this study.** Highlighted are restriction enzyme recognition sites for SfiI (blue) and BsaI (yellow), and the *ccdB* operon (green). BsaI digest overhangs are underlined. Nucleotides targeted for mutagenesis are lowercase and bold.

| | |
|---|---|
| Parts 1; 6 with ccdB operon | GGCCCAGCCGGCCATGGCTCAAGTGCAGCTGGTAGAATCTGGTGGCGGTCTGGTTCAGGCTGGCGGTTCCCTGCGTCTGTCCTGTGAGAGACCGAATACATACATGTCGTAATACGACTCACTATAGGGGCACGCGTCTTCTGCGGCCGCATTAGGCACCCCAGGCTTGACAGCTAGCTCAGTCCTAGGTATAATGCTAGCCCTTGTGTTGTTATCCGCTCACAAAAAGAGGAGAAAGGATCCATGCAGTTTAAGGTTTACACCTATAAAAGAGAGAGCCGTTATCGTCTGTTTGTGGATGTACAGAGTGATATTATTGACACGCCGGGGCGACGGATGGTGATCCCCCTGGCCAGTGCACGTCTGCTGTCAGATAAAGTCTCCCGTGAACTTTACCCGGTGGTGCATATCGGGGATGAAAGCTGGCGCATGATGACCACCGATATGGCCAGTGTGCCGGTGTCTGTTATCGGGGAAGAAGTGGCTGATCTCAGCCACCGCGAAAATGACATCAAAAACGCCATTAACCTGATGTTCTGGGGAATATAATAACGCAAAAAACCCCGCCCCTGACAGGGCGGGGTTTTTTCGCCAGAAGACCTCCATGCAGAGTCGTACTAGGTCTCTACCGTATCTGCTGGTAGGGCTGGTCACCACCATCACCATCACGAACCTGAAGCCCAGTACCCGTACGACGTTCCGGACTACGGTTCCGCGGCCTCGGGGGCC |
| Part 2; CDR1 | GGTCTCATGTGCCGCAAGC**ggtttcccg**GTGt**actggaactct**ATG**tac**TGG**tatcgtcag**GCACCGGGCAAGGAATGAGACC |
| Part 3; CDR2 | GGTCTCAGGAACGTGAGTGGGTCGCG**gcgattactagccagggtggtggtacggaatacgcagat**TCTGTGATGAGACC |
| Part 4 | GGTCTCAGTGAAGGGCCGCTTTACCATCAGCCGCGACAACGCGAAGAATACGGTCTATTTGCAGATGAATAGCCTGAAACCGGAAGATACCGCGGTTTACTACTGAGACC |
| Part 5; CDR3 | GGTCTCACTACTGT**act**GTG**ggtgtgggtgtttcttaccgtggccaaggtacc**CAAGTGACCGTGAGACC |

### Supplementary Figures

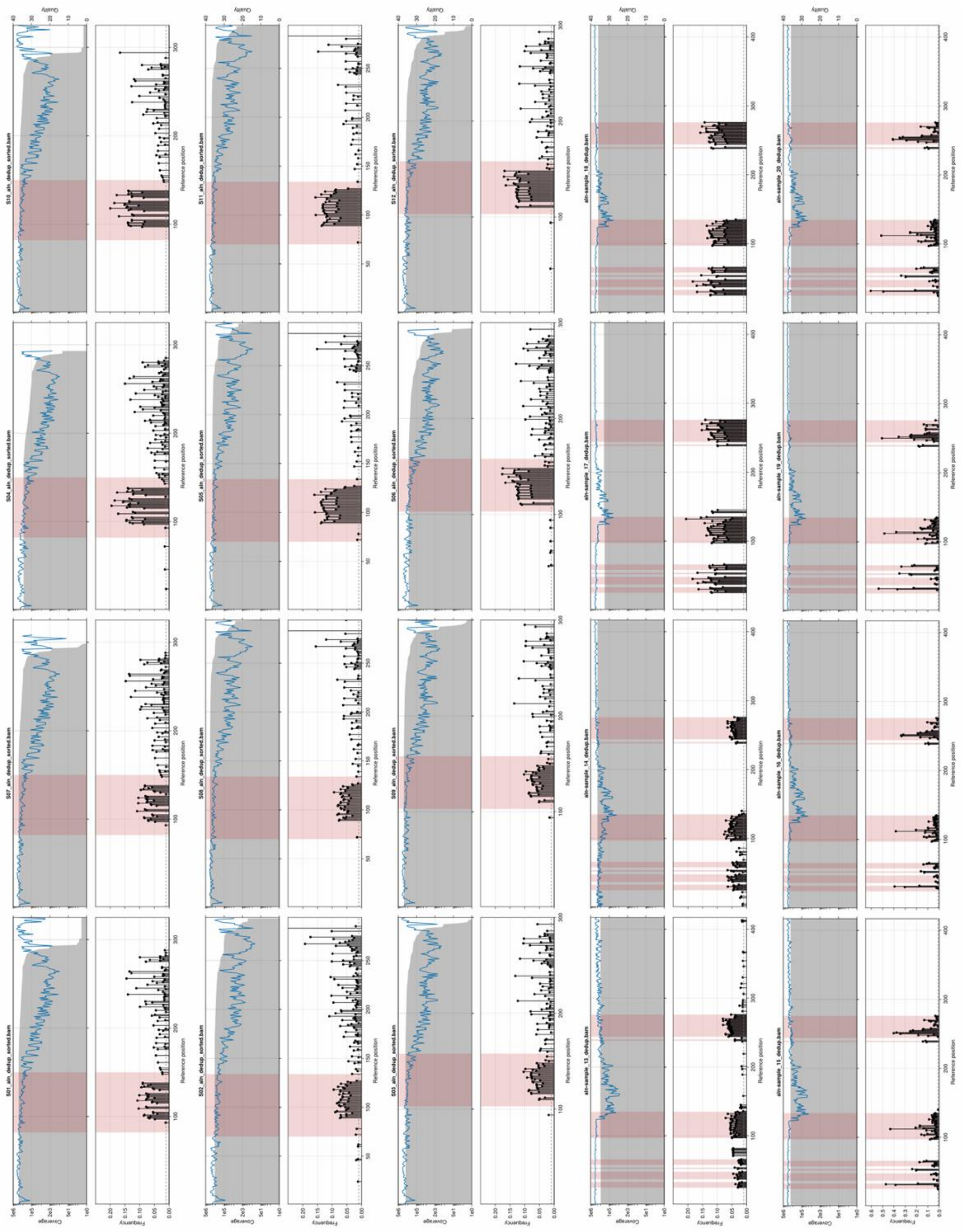

**Figure S1.** Alignment of deduplicated (post-QC; pre-correction) reads. Coverage (grey fill), quality (blue line) across the reference positions, with mismatches (black stems) displayed per position where frequency > threshold (0.1). No insertions or deletions observed > threshold. Highlighted in red are the duplexed oligo part (S01-12), or targeted residues (S13-20). S01, 07 CDR1 single; S02, 08 CDR2 single; S03, 09 CDR3 single; S04, 10 CDR1 dual; S05, 11 CDR2 dual, S06, 12 CDR3 dual; S13, 14 Dual DMS Pre; S15, 16 Dual DMS post; S17, 18 CMB pre; S19, 20 CMB post.

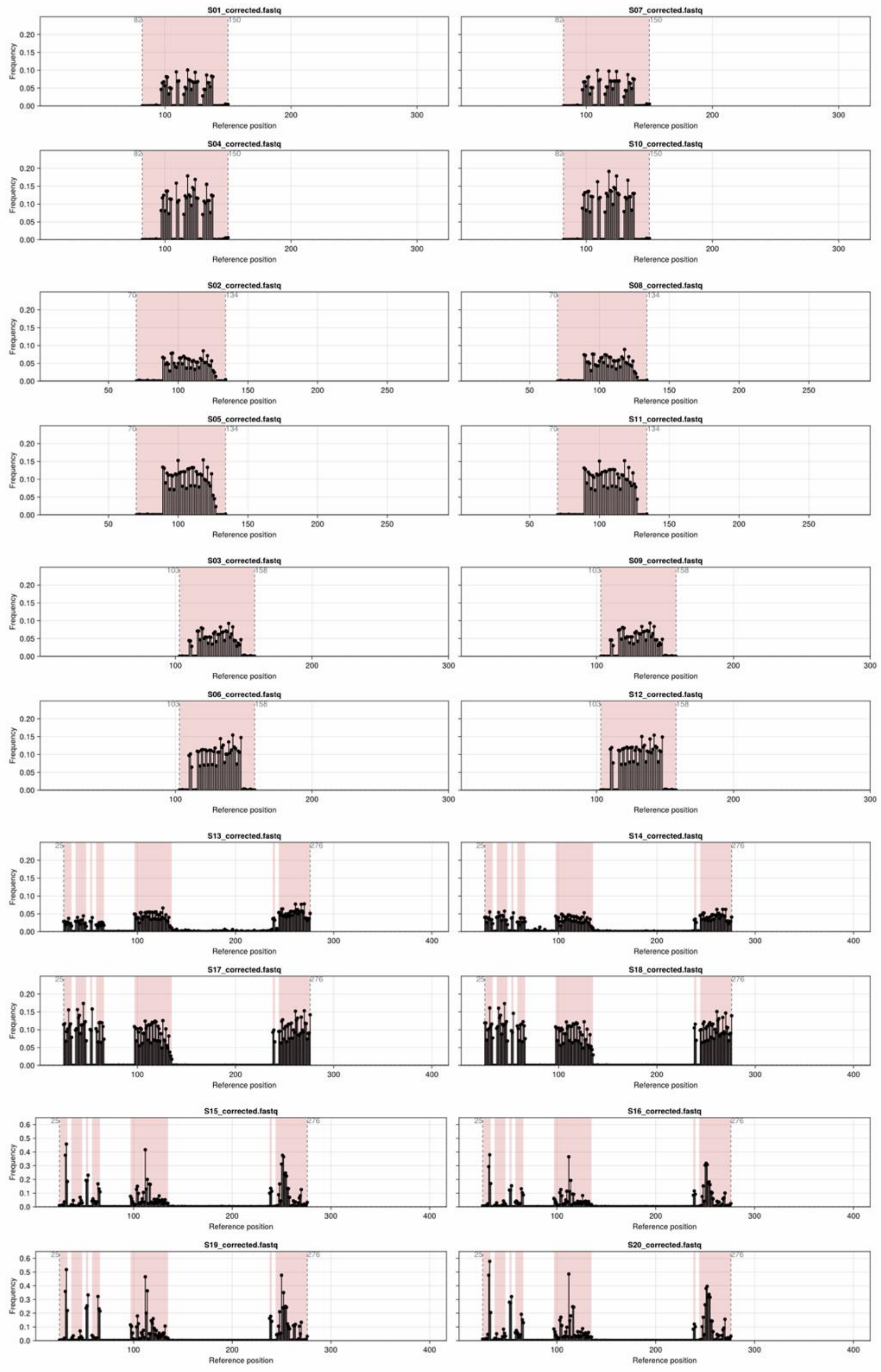


**Figure S2.** Alignment of corrected reads (Q<20 reverted to WT nt). Mismatches (black stems) displayed per position. Highlighted in red are the duplexed oligo part (S01-12), or targeted residues (S13-20). Read trimming boundaries are also shown (grey dashed line, text). S01, 07 CDR1 single; S02, 08 CDR2 single; S03, 09 CDR3 single; S04, 10 CDR1 dual; S05, 11 CDR2 dual, S06, 12 CDR3 dual; S13, 14 Dual DMS Pre; S15, 16 Dual DMS post; S17, 18 CMB pre; S19, 20 CMB post.
